# Non-canonical histone H3.3 and its chaperones HIRA and DAXX participate in the regulation of KSHV latency

**DOI:** 10.64898/2026.04.02.716100

**Authors:** Sarah McMahon, Vaibhav Jain, Viacheslav Morozov, Sunantha Sethuraman, Jianhong Hu, Ritu Shekhar, Netanya Keil, Peter Turner, Alexander Ishov, Rolf Renne

## Abstract

Kaposi’s sarcoma-associated herpesvirus (KSHV), also named HHV-8, is the etiological agent of Kaposi sarcoma (KS), Primary effusion lymphoma (PEL), and Multicentric Castleman’s disease. After de novo infection, KSHV genomes rapidly circularize and acquire a chromatin state that favors latency. During latency, the KSHV episome is decorated with distinct epigenetic marks that segregate the viral genome into transcriptionally active and repressed domains, enabling persistent silencing of lytic genes while retaining the capacity for reactivation. Transcription activity of chromatin is regulated at multiple levels, including the incorporation of histone variants such as H3.3, by a specific set of histone chaperones such as HIRA and DAXX. The interaction between LANA and these interphase active chaperones suggests that H3.3 deposition is a critical driver of early chromatinization and the long-term stability of KSHV latency. We detected rapid H3.3 deposition on KSHV episomes and on episomes within long-term infected cells. Moreover, we demonstrated that genetically disrupting the host H3.3 chaperone HIRA pathway by CRISPR/Cas9-mediated knockout impacted the regulation of LANA and maintenance of viral latency that was not altered in DAXX knockout cells. Collectively, these results support a role for HIRA-mediated H3.3 deposition in the regulation of KSHV latency.

## Introduction

Kaposi’s sarcoma-associated herpesvirus (KSHV) is the causative agent of several human cancers including Kaposi sarcoma (KS)^1^ and the lymphoproliferative diseases: Primary effusion lymphoma (PEL)^2^ and Multicentric Castleman’s Disease (MCD)^3^. A characteristic feature of KSHV pathogenesis is the predominance of a latently infected cell state where long-term persistence supports tumorigenesis. Upon infection of host cells, the KSHV genome is rapidly packaged into chromatin (episomes), utilizing host chromatin machinery that partitions the genome into transcriptionally active and repressed domains^4^. This process is essential for sustained expression of a subset of latency genes, including the Latency Associated Nuclear Antigen (LANA). Upon establishment of viral latency, replication and segregation of viral genomes is essential for long-term viral persistence.

The regulation of latency expression is dictated by several processes, including modulation of host histone^5^. Eukaryotic cells express three types of histone H3 variants: H3.1, H3.2, and H3.3^6^. Histone H3.1 and H3.2 are the predominant (“canonical”) variants that are expressed solely during S phase of the cell cycle and are exclusively incorporated into newly synthesized DNA^7–9^. The variant histone H3.3 differs from H3.1 by only five amino acids, and is highly conserved in mammals^10^. Comparable to canonical H3.1 and H3.2, H3.3 histone tails also undergo epigenetic and biochemical changes^11, 12^. However, unlike the canonical H3.1, histone H3.3 is expressed throughout the cell cycle, can be assembled into chromatin outside of S-phase^13^, and is mostly enriched in regions with elevated transcriptional activity^14^. In accord with their availability throughout the cell cycle, histone H3.3 is deposited by unique chaperone complexes, including the histone cell cycle regulator A (HIRA) and death-domain associated protein (DAXX) complexes. HIRA is responsible for deposition of H3.3 into regions of transcriptional activity, nucleosome-free regions, and at sites of DNA damage ^14–17^. Conversely, DAXX, a PML Nuclear Bodies/ND10-associated protein, is more commonly associated with transcriptionally repressed heterochromatin where it deposits H3.3 to specific transcription factor binding sites, pericentromeric regions, and telomeres ^16, 18–20^.

Evidence suggests that large double-stranded DNA viruses, such as KSHV and EBV, utilize alternative histones and their chaperones during *de novo* viral infection^21^. For EBV, it has been shown that the tegument protein BNRF1 suppresses ATRX-DAXX-mediated H3.3 loading onto viral episomes early after infection^22^. KSHV LANA interacts with HIRA, DAXX, and DEK, which all function as H3.3 chaperones^23–25^. LANA also interacts with core components of the HIRA complex and with downstream effectors including the FACT chromatin remodeling complex^25, 26^. Moreover, it has been shown that the recruitment of the Polycomb Repressive Complex-2 (PRC2) complex on epigenetically bivalent and transcriptionally poised promoters is mediated by HIRA-dependent incorporation of H3.3^27^. DAXX was also identified from a proteomic screen as an interactor of LANA^25^, and this interaction was recently demonstrated to be critical for epigenetic maintenance of episomes associated with LANA nuclear bodies during KSHV latency^28^. However, whether the enrichment of alternate histone H3.3, mediated by the chaperones HIRA and DAXX, is involved in the establishment and maintenance of latent KSHV episomes has not been established.

Here, we tested the hypothesis that the histone variant H3.3 contributes to the establishment and maintenance of KSHV latency. We report that H3.3 is enriched on the latent KSHV genome in PEL BCBL-1 cells and latently infected HEp-2 cells and is deposited on the viral episome during early stages of latency establishment. We further show that the H3.3 histone chaperones HIRA and DAXX colocalize with LANA on KSHV episomes independently of PML/ND10. Using knockout cell lines, we demonstrate that HIRA, but not DAXX, regulates H3.3 positioning within the major viral latency region encompassing LANA. Together, these findings support a role for the H3.3 pathway, particularly HIRA-mediated activity, in the establishment and maintenance of KSHV latency.

## Results

### Histone H3.3 is deposited during the pre-latency phase after *de novo* infection

During *de novo* infection, KSHV releases naked linear DNA into the nucleus of host cells where it circularizes to form viral episomes. The chromatinization of KSHV episomes has been demonstrated as early as two hours post-infection^19, 29^. To test whether H3.3 contributes to early chromatinization, we infected SLK cells with recombinant KSHV/BAC16 and examined H3.3 deposition by ChIP-qPCR on multiple viral promoters at 2, 8, 24, 48, and 72 hrs post infection (Figure 1A). H3.3 deposition was detectable on the RTA promoter and late gene ORF19 promoter at 2 hrs post infection, whereas LANA showed no H3.3 enrichment at this earliest timepoint. At 8 and 24 hrs post infection, H3.3 was detected on the RTA promoter and early gene K7 promoter, which are known to harbor bivalent marks, and to a lesser extent the ORF19 promoter. At late time points (48 and 72 hrs), H3.3 association with the LANA promoter increased while its association with the lytic promoter declined. These data are consistent with early work by others showing that both the RTA and LANA genes are transcribed very early, but that the RTA gene is subsequently silenced in response to increased LANA expression^30^. Collectively, these data indicate that H3.3 incorporation is an early and dynamic feature of KSHV episome chromatinization that supports latency establishment.

**Figure 1.**
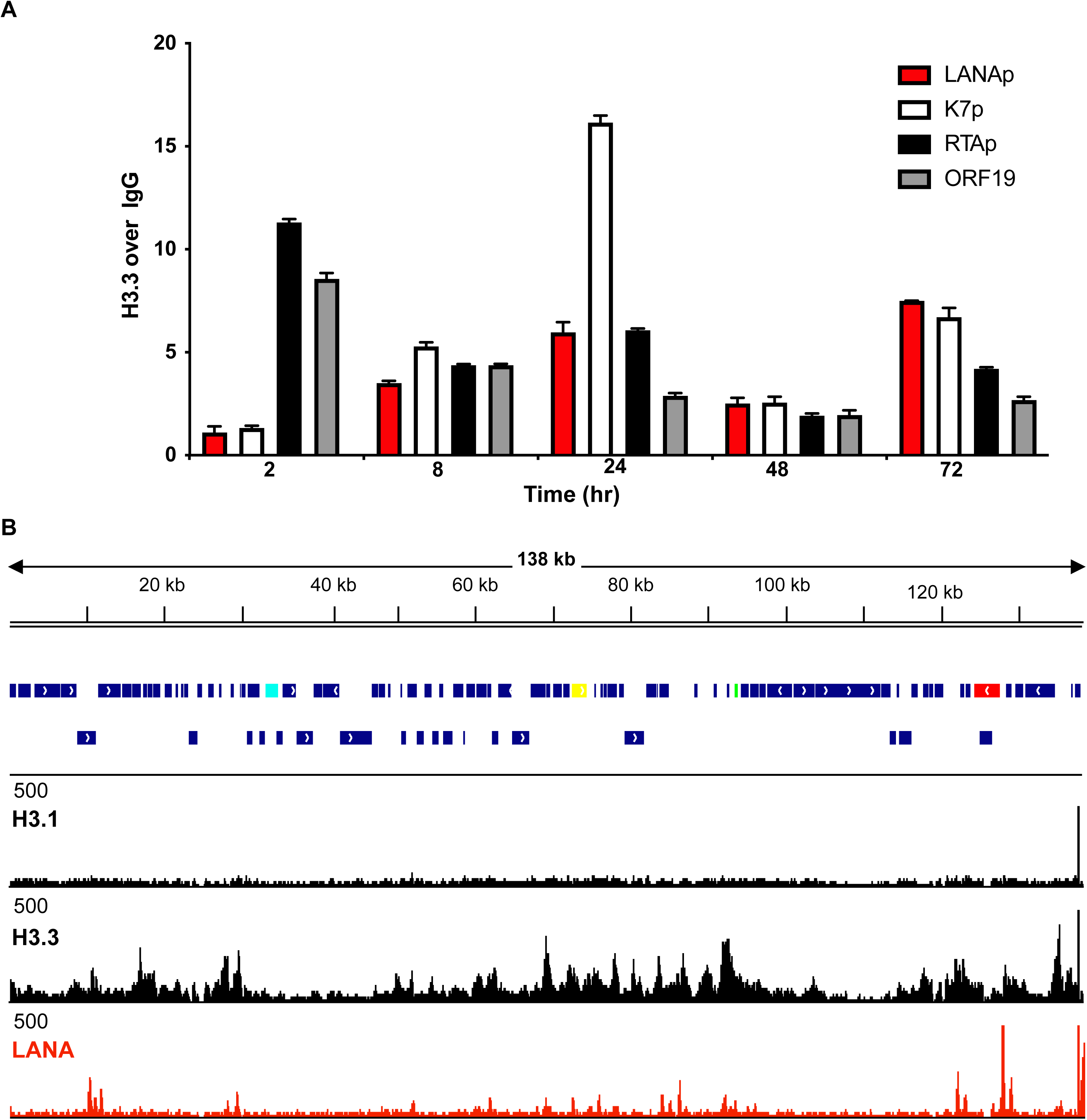
Histone H3.3 is incorporated in the KSHV episome early after de novo infection and in latently infected PEL cells. (A) Kinetic analysis of H3.3 deposition onto latent and lytic promoters after de novo infection. Histone H3.3 occupancy was determined by ChIP-qPCR at the indicated time points after *de novo* infection of SLK cells with KSHV/BAC16. H3.3 ChIP was performed using anti-Histone H3.3 (phospho S31) antibody. Occupancy of the LANA, K7, and RTA promoters, and ORF19 gene body is shown. Enrichment is expressed as fold increase over IgG control. (B) ChIP-seq assays were performed in BCBL-1 cells stably expressing HA-tagged canonical histone H3.1 (HA-H3.1) or HA-tagged histone variant H3.3 (HA-H3.3) using an anti-HA antibody to detect tagged histone. LANA ChIP-seq was performed in BCBL-1 cells with LANA monoclonal antibody. The ChIP-seq peaks for HA-tagged H3.1, H3.3, and LANA are shown aligned to the KSHV genome. The genome track shows KSHV genome organization with genes of interest highlighted latent (LANA – red), immediate early (RTA - yellow), early (K11 - green), and late (ORF19 – cyan).

### H3.3 is present on latent viral episomes

After establishing H3.3 deposition and its chaperones are active during KSHV infection of SLK cells, we wanted to test whether H3.3 is also present on KSHV episomes in infected lymphoid cells. H3.1 and H3.3 only differ by five amino acids, therefore making it difficult to raise an antibody that specifically differentiates these two histone molecules. Therefore, to investigate the presence of histone H3.3 on the KSHV episome, we engineered patient-derived BCBL-1 cells to ectopically express either HA-tagged canonical H3.1 or H3.3 by stable lentivirus transduction and verified HA-H3.1 and HA-H3.3 expression by western blot (Supplementary Figure 1). We performed ChIP-seq to map ectopically expressed H3.1 and H3.3 (anti-HA) and LANA to examine their distributions across the KSHV episome (Figure 1B). LANA is enriched on several viral loci, including the KSHV latency-associated region (KLAR), viral Interferon Regulatory Factors (vIRFs), and the lytic switch gene RTA. Canonical histone H3.1 was detected broadly and relatively uniformly throughout the viral genome, whereas histone H3.3 localized as discrete peaks. Except for a cluster of H3.3 peaks spanning the ORF11 to ORF2 region (16.5 kb-18.7 kb), the majority of H3.3 peaks colocalize with LANA binding sites. This co-occupancy of H3.3 and LANA and the association between LANA and the H3.3 chaperone pathways suggest a possible role for LANA in H3.3 deposition. To validate these findings on endogenous histones of infected cells, we performed ChIP-qPCR in parental BCBL-1 cells using a H3.3-S31P-specific antibody. The promoters of LANA, K7, and the ORF19 gene were all enriched for endogenous H3.3 (Supplementary Figure 1). Together these data demonstrate that latent KSHV episomes are decorated with H3.3 at multiple loci and H3.3 and LANA show co-occupancy on the viral episome.

### H3.3 chaperones HIRA and DAXX interact with and colocalize to LANA speckles in KSHV infected cells

Previous studies have identified LANA to be associated with the H3.3 chaperones HIRA and DAXX^25, 28, 31^. Both chaperones are known to be involved in host immune response to foreign DNA^21^ and in remodeling PML/ND10 nuclear bodies^32–38^. KSHV/LANA form unique nuclear speckles to support viral latency and given the interaction of LANA with H3.3 histone chaperones we asked whether DAXX and HIRA affect LANA speckle organization. First, we validated the interaction of KSHV LANA with DAXX and HIRA by coimmunoprecipitation with exogenous overexpression of LANA with either DAXX or HIRA. This demonstrated that both DAXX and HIRA interact with LANA (Figure 2A). To determine whether these interactions reflect spatial proximity in infected cells, we performed immunofluorescence assays (IFA) to compare the localization of LANA, HIRA, DAXX, and PML/ND10 components (Figure 2B). In mock-infected cells, DAXX localized to discrete nuclear bodies that overlapped extensively with PML/ND10 (Figures 2Bi-2Biv), whereas HIRA displayed a diffuse nuclear distribution with only limited association with PML bodies (Figures 2Bv-2Bviii). Cells infected with KSHV displayed characteristic LANA nuclear speckles^39^ (Figures 2Bix and 2Bxiii) that are spatially distinct from PML/ND10^19^. Interestingly, in KSHV-infected cells, DAXX colocalizes with PML as in uninfected cells, while a portion of DAXX is redistributed and colocalizes with LANA, outside of PML/ND10 (Figures 2Bix – 2Bxii). HIRA also showed pronounced colocalization with LANA speckles in infected cells, independent of PML/ND10 (Figures 2Bxiii – 2Bxvi). Overall, these data demonstrate that in KSHV infected cells both H3.3 chaperones DAXX and HIRA accumulate with KSHV LANA outside of PML/ND10 and that KSHV infection does not affect PML/ND10 structures.

**Figure 2.**
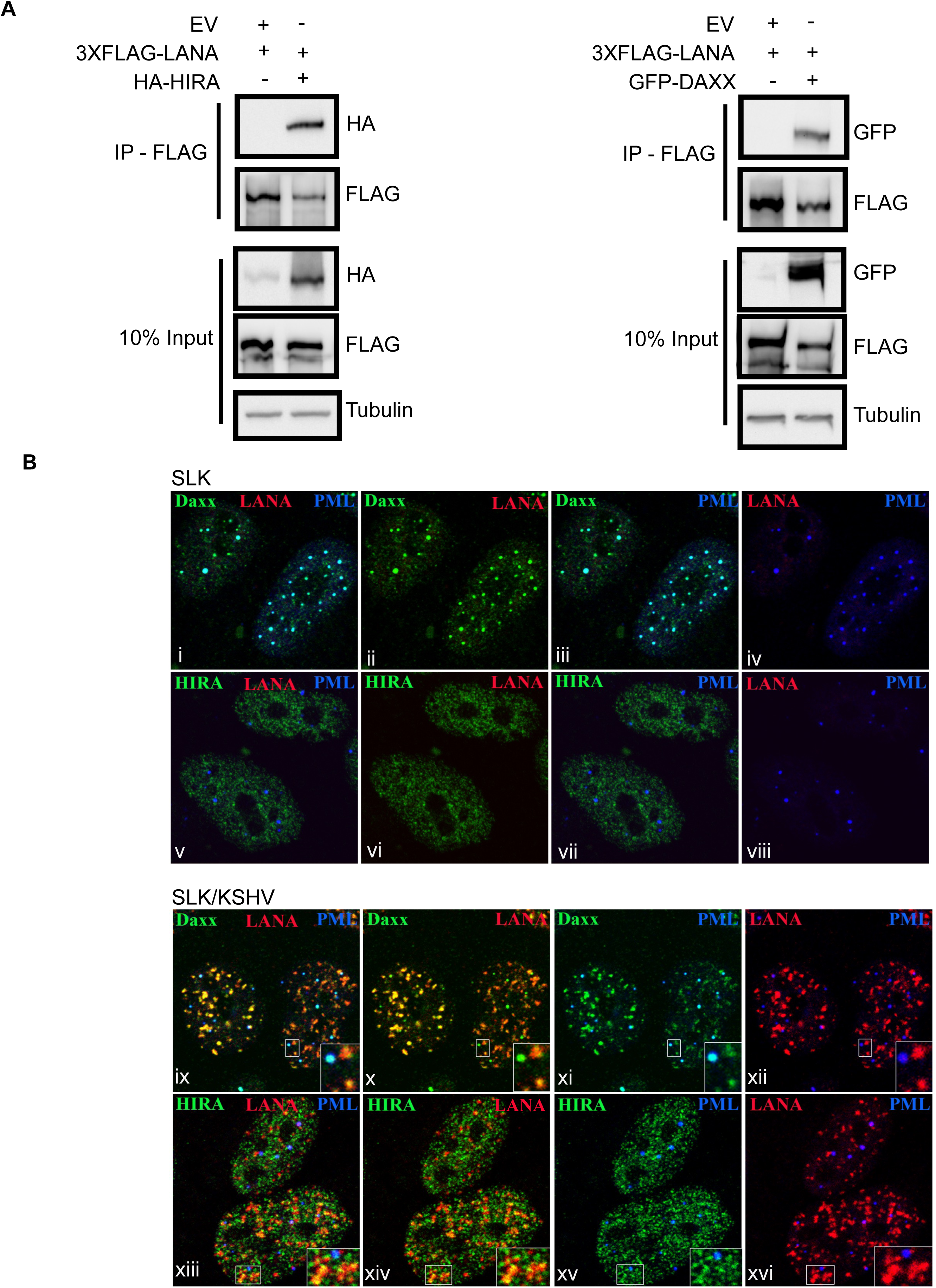
H3.3 chaperones DAXX and HIRA interact with LANA and colocalize to LANA speckles in KSHV-infected SLK cells. (A) Interaction of LANA with histone chaperones by coimmunoprecipitation of FLAG-LANA with HA-HIRA and GFP-DAXX in 293T cells. (B) Top two rows (i - viii): uninfected SLK cells; bottom two rows (ix - xvi): KSHV-infected SLK cells. (i – iv): Localization of DAXX (green) and ND10/PML (blue). (v - viii) HIRA localization (green) in uninfected SLK cells. DAXX colocalizes with ND10/PML NBs (PML), while HIRA has minor accumulation at ND10/PML NBs. Third row (infected SLK cells, ix - xii): DAXX (green) colocalizes with LANA (red) and is associated with ND10/PML NBs (PML, blue). The inset at the bottom right demonstrates the colocalization of DAXX with both LANA and PML. Bottom row (xiii - xvi): HIRA (green) has substantial colocalization with LANA (red), and minor accumulation at ND10/PML NBs (PML, blue).

Given the association of LANA with H3.3 loading on viral genomes and colocalization of LANA with the histone chaperones DAXX and HIRA, we were interested in dissecting the genetic relationship between LANA and these H3.3 chaperones. A panel of LANA domain constructs previously described^40^ were used for coimmunoprecipitation experiments with epitope tagged HIRA and DAXX constructs (Supplementary Figure 2A). No specific domain was identified as required for binding, instead interactions with both HIRA and DAXX were strongest when the large, intrinsically disordered region of LANA was present. These results posture that LANA-chaperone association is mediated by multivalent contacts characteristic of proteins that participate in bimolecular condensates^41^.

### CRISPR/Cas9-based knockout of H3.3 chaperones does not affect LANA speckles in latently infected cells

Given widespread H3.3 loading across the KSHV genome and the interaction and colocalization of DAXX and HIRA with LANA speckles, we next tested the genetic contribution of each H3.3 chaperone pathway to viral latency maintenance. Using CRISPR/Cas9-based gene editing^42, 43^ in HEp-2 larynx carcinoma cells we generated DAXX and HIRA knockout (KO) cell lines and introduced a FLAG tag into one of the endogenous H3.3 genes (H3FB) to circumvent specificity limitations of commercially available H3.3 antibodies. Clones were validated by western blot for expression of DAXX, HIRA, and FLAG-H3.3, confirming expression of the knock-in FLAG-H3.3 allele and lack of expression of DAXX and HIRA in the appropriate KO cells (Supplementary Figure 3A). For downstream experiments, one clone of CRISPR control and two clones of each of the HIRA KO and DAXX KO were used. The FLAG-H3.3 knock-in was characterized by IFA and metaphase spreads, confirming chromosome association of FLAG-H3.3 (Supplementary Figure 3B top and middle rows). Lastly, in KSHV infected cells FLAG-H3.3 was diffuse throughout nuclei with intact nuclear LANA speckles (Supplementary Figure 3B bottom row).

To examine the localization of DAXX, HIRA, and LANA in these KO KSHV infected lines, we performed IFA in these HEp-2 lines (Figure 3). In control KSHV infected cells, DAXX closely associated with LANA speckles (Figure 3A-D), and HIRA colocalized with LANA (Figure 3E-H) consistent with observations in iSLK cells (Figure 2). In HIRA KO cells, HIRA signal is absent and does not affect DAXX/LANA colocalization (Figure 3I-L). However, it is interesting to note that compared to control cells where DAXX is highly localized in LANA speckles (Figure 3B), in HIRA KO cells while some LANA-DAXX interactions are maintained, DAXX is more diffusely distributed within the cell independent of LANA (Figure 3J). Previous studies have shown that upon lytic induction of latently infected cells that DAXX is evicted from LANA speckles^28^, perhaps suggesting that HIRA KO cells might support viral episomes more permissive to a lytic state. Additionally, neither PML/ND10 or LANA speckles are affected by HIRA KO (Figure 3M-P). In KSHV infected DAXX KO cells, DAXX staining is absent (Figure 3Q-T), and no changes in HIRA association with or formation of PML/ND10 or LANA speckles was identified (Figure 3U-X). Overall, the loss of one of the H3.3 chaperones did not affect the colocalization of the remaining chaperone with LANA and did not perturb PML/ND10 architecture.

**Figure 3.**
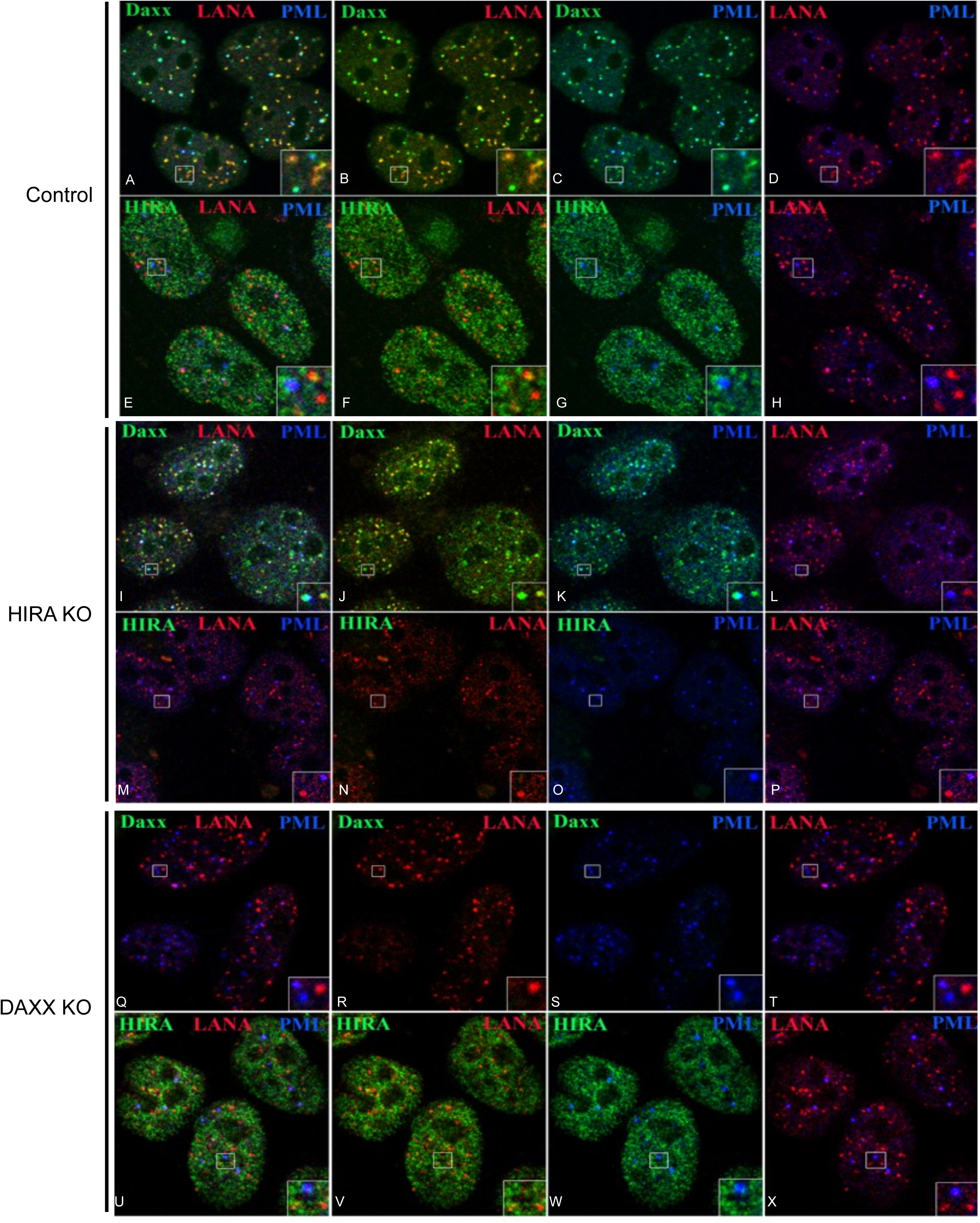
Characterization of DAXX KO and HIRA KO in KSHV latently infected cells. (A - H) Distribution of H3.3 chaperones DAXX and HIRA in KSHV infected HEp-2 cells. Top row (A - D): DAXX (green) colocalizes with LANA (red) and is associated with ND10/PML NBs (PML, blue) in four representative nuclei. The inset at the bottom right demonstrates the colocalization of DAXX with both LANA and PML. Second row (E - H): HIRA (green) has minor colocalization with LANA (red), and ND10/PML NBs (PML, blue). I - P shows the distribution of DAXX and HIRA in KSHV infected HIRA KO HEp-2 cells. (I - L): DAXX (green) colocalizes with LANA (red) and is associated with ND10/PML NBs (PML, blue), similar to control HEp-2 cells (in A - D above). HIRA KO does not affect the intranuclear distribution of DAXX. (M – P): No visible HIRA (green); LANA (red), and ND10/PML NBs (PML, blue) are not affected by HIRA KO. (N) Increased diffuse signal of DAXX (green) compared to control (B). Q - X show distribution of DAXX and HIRA in KSHV infected DAXX KO HEp-2 cells. (Q - T): No visible DAXX (green); LANA (red), and ND10/PML NBs (PML, blue) are not affected by DAXX KO. (U – X): HIRA (green) has minor colocalization with LANA (red), and ND10/PML NBs (PML, blue), like control HEp-2 cells (A). DAXX KO does not affect the intranuclear distribution of HIRA.

### HIRA and DAXX are both utilized to load H3.3 onto KSHV episomes

HIRA and DAXX deposit histone H3.3 into transcriptionally active and repressive chromatin, respectively, but their roles during KSHV latency are not well defined. To map the HIRA- and DAXX-dependent histone H3.3 deposition on latent viral genomes, we performed CUT&RUN in latently infected cells^44^. First, we compared H3.3 distribution on the KSHV episome between PEL BCBL-1 cells and HEp-2 latently infected cells. Overall, both cell types show similar, ubiquitous H3.3 loading across the viral genome (Figure 4A). Minor differences in peak patterns were observed at a few loci that may reflect biologically meaningful variation between cell types or arise due to technical differences between ChIP-seq and CUT&RUN or from ectopic versus endogenous H3.3 expression.

**Figure 4.**
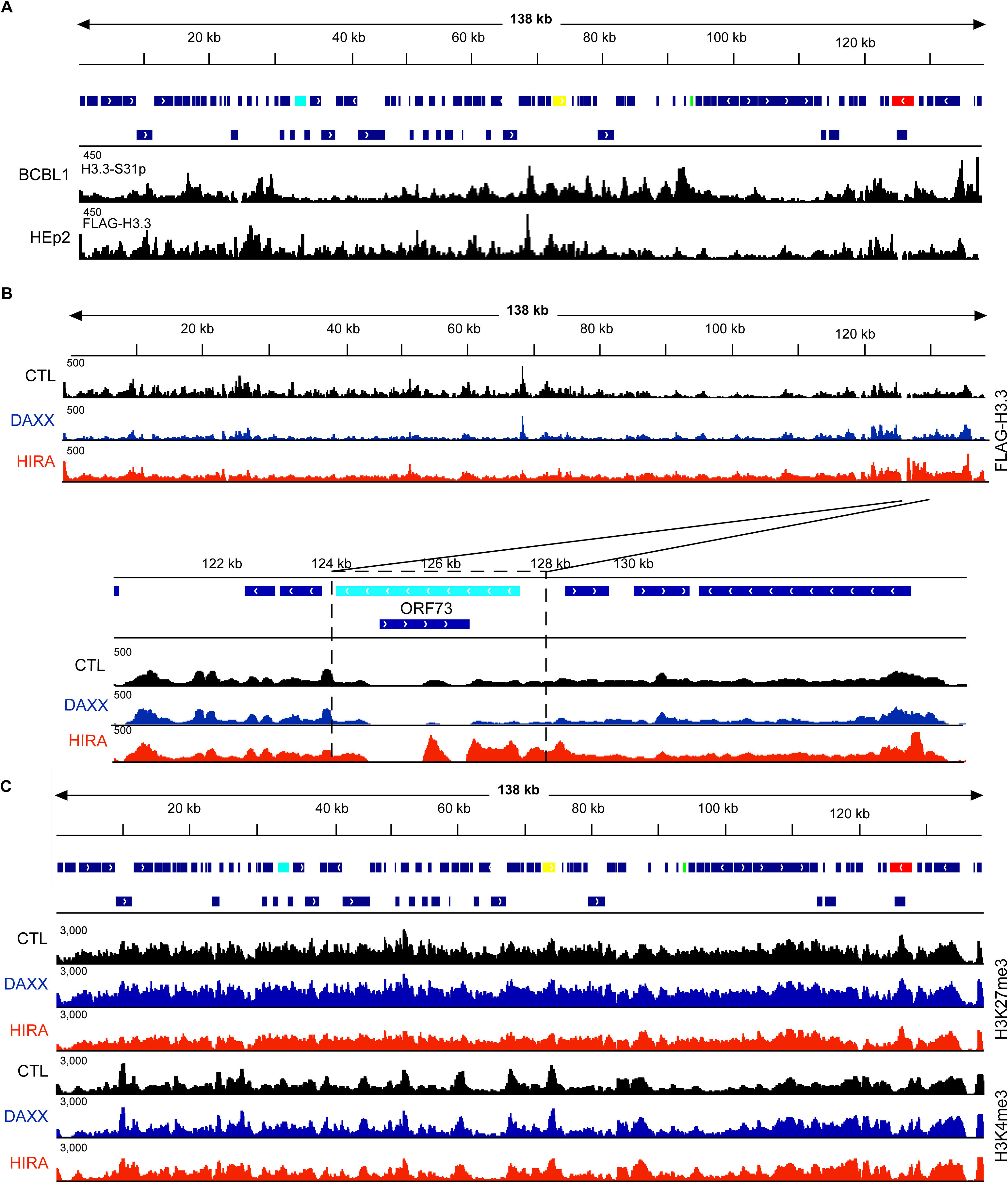
Histone chaperones DAXX and HIRA both load H3.3 onto KSHV episomes. (A) H3.3 loading across KSHV episome in BCBL1 (ChIP-seq as depicted in Figure 1) and FLAG-tagged H3.3 in HEp-2 cells (CUT&RUN). (B) FLAG H3.3 CUT&RUN in wild-type control, DAXX KO, and HIRA KO HEp-2 KSHV infected cells. Lower panel is zoomed in on ORF73 (LANA) locus. (C) ChIP-seq for histone modifications (H3K27me3 and H3K4me3) in wild-type control, DAXX KO, and HIRA KO across viral episomes in HEp-2 KSHV infected cells. For each panel, the genome track shows KSHV genome organization with the following viral latent (LANA – red), immediate early (RTA - yellow), early (K11 - green), and late (ORF19 – cyan) genes highlighted for reference.

We next wanted to determine whether H3.3 chaperone deletion could alter H3.3 loading onto the viral episome. We compared CUT&RUN from control, HIRA KO, and DAXX KO cells with an antibody to FLAG-tagged H3.3 (Figure 4B). During data processing, we detected selection biases for smaller episomes in some samples (Supplementary Figure 4A), a phenomenon reported previously in infected cells^24, 45^. Therefore, to overcome this artifact we examined the signal across sequenced input DNA. From this we identified a large region on the left side of genomes that was particularly reduced in control and DAXX KO samples. Therefore, a scale factor was generated for all samples to scale the left side of genomes to right (Supplementary Figure 4B).

After adjustment, we compared FLAG-H3.3 loading between control, HIRA KO, and DAXX KO cells (Figure 4B). Overall, we observed H3.3 loading all along the viral genome independent of genotype. Unexpectedly, we identified that in the region coding for LANA there was an increased signal unique to HIRA KO cells when compared to control and DAXX KO (Figure 4B, zoom in). In addition to H3.3 loading, we also profiled active (H3K4me3) and repressive (H3K37me3) histone modifications across control and KO samples to see if whether these histone chaperones regulate the partitioning the episome into transcriptionally active and repressive regions (Figure 4C). Looking at these marks across the viral genome demonstrated there was no major changes to these histone modifications suggesting the impact of these H3.3 chaperones on the viral episome is more important for loading and less on partitioning the genome into transcriptionally active and repressive regions via epigenetic modification of histones. Overall, our findings indicate both HIRA- and DAXX-mediate incorporation of H3.3 onto viral genomes, with exception of the LANA locus that is affected by HIRA KO.

### HIRA loss increases H3.3 at the LANA locus and derepresses viral gene expression

The major finding in our characterization of DAXX- and HIRA-dependent loading of H3.3 was a selective increase in H3.3 occupancy across the LANA locus in HIRA KO cells. To test whether this altered H3.3 loading affects viral transcription, we measured expression of LANA, RTA, and ORF19 by RT-qPCR in latently infected control, DAXX KO, and HIRA KO cells. In the uninduced latent state, HIRA KO cells, but not DAXX KO cells, exhibited significantly elevated RTA and ORF19 transcript levels compared to control (Figure 5A). Under sub-optimal induction conditions that are insufficient to drive robust lytic activation, this increased effect is lost, suggesting that loss of HIRA is critical for maintenance of viral latency rather than productive lytic induction. Consistent with RT-qPCR data, infection with the reporter .219 KSHV recombinant virus revealed detectable PAN-promoter driven RFP in uninduced HIRA KO cells (∼9 RFP^+^ cells per field), a signal that did not appreciably increase under sub-optimal induction (∼7 RFP^+^ cells per field) (Figure 5B). To assess whether these transcriptional changes translate into infectious output, we performed supernatant transfer from uninduced cells onto naïve 293T cells and quantified GFP^+^ recipient cells by flow cytometry at 48 hours. Supernatant from control cells produced minimal GFP^+^ infection (0.24%), while DAXX KO supernatant showed a modest increase (0.675), and HIRA KO supernatant produced a larger rise (1.01%) (Figure 5C). Together, these results indicate that loss of HIRA alters H3.3 positioning at the LANA locus and weakens latency control, leading to increased expression of lytic transcripts and a modest rise in virus release, without triggering widespread lytic replication.

**Figure 5.**
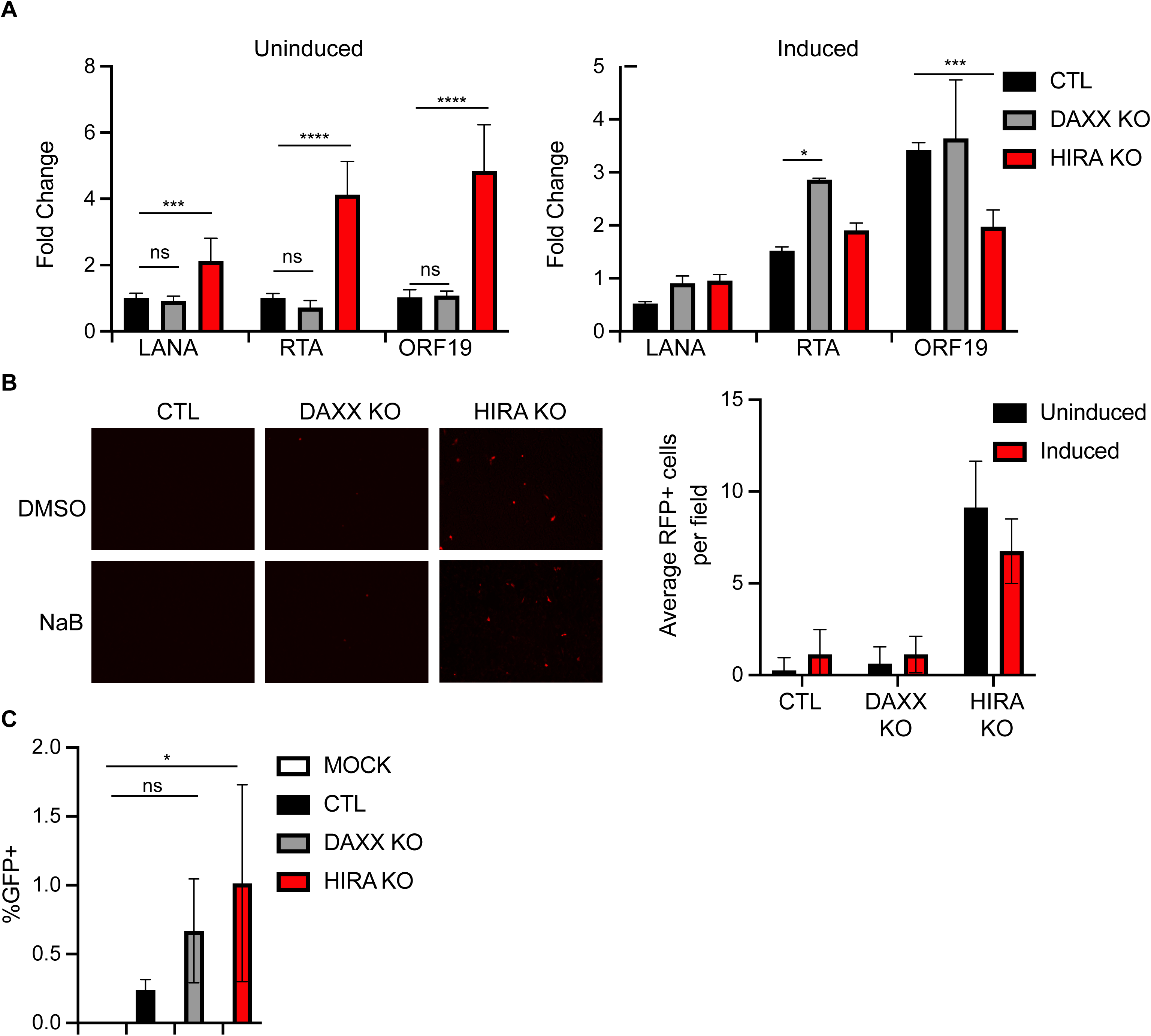
HIRA KO disrupts maintenance of viral latency in HEp-2 KSHV infected cells. (A) Wild-type control (CTL), DAXX KO, and HIRA KO HEp-2 cells were infected with KSHV/BAC16 virus, and latently infected cells were selected using hygromycin. Infected cells were either uninduced (panel A; left) or sub-optimally induced with 1 mM sodium butyrate (panel A; right) for 48 hours. Transcripts of LANA, RTA, and ORF19 were quantified by RT-qPCR normalized to GAPDH. In uninduced cells, data is represented as relative to CTL. For sub-optimal induction experiments data is represented as relative to respective uninduced expression. Error bars indicate the SD of three biological replicates and significance of p-values are represented by asterisks (*p<0.1, ** p<0.01, ***p<0.001, ****p<0.0001). (B) Wild-type control (CTL), DAXX KO, and HIRA KO HEp-2 cells were infected with KSHV/.219 dual reporter virus. RFP representing lytic activation was examined in uninduced or sub-optimally induced cells and presented as fluorescence images (left) and quantified across multiple fields (right). (C) Supernatant from uninduced control (CTL), DAXX KO, and HIRA KO HEp-2 cells was used to transduce naïve 293T cells and GFP positivity was determined at 24 hours post transduction by flow cytometry.

## Discussion

During latent infection, KSHV acquires and maintains a distinct chromatin state by utilizing host epigenetic machinery. Our work provides additional evidence of direct involvement of the non-canonical histone variant H3.3 and its chaperones HIRA and DAXX in the establishment and maintenance of the latent KSHV episome. We show that HIRA and DAXX H3.3 histone chaperones interact with and colocalize with KSHV-LANA in KSHV-infected cells independent of PML/ND10. We also observed that loss of HIRA, but not DAXX, alters H3.3 deposition onto the KSHV episome, altering loading at the major latency locus LANA with consequences on viral gene regulation and latency.

Contrary to the canonical histone H3.1 and H3.2 that are deposited solely during DNA replication in S-phase, deposition of histone variant H3.3 occurs throughout interphase and is presumably available during viral infection in a cell-cycle independent manner. Therefore, we hypothesized that deposition of histone H3.3 plays an important role in early steps of latency establishment, since during KSHV infection LANA interacts either directly with both HIRA and DAXX or indirectly with other downstream components of the HIRA pathway such as Spt16^19, 25, 26, 31^. We observed that H3.3 is deposited on KSHV episomes as early as 2 hours after *de novo* infection prior to the establishment of latency and is present during long-term latency. This suggests that the H3.3 pathway is involved early in the process of KSHV latency establishment. However, future analyses using these genetic knockout lines will be important to determine the relative contribution of HIRA and DAXX on H3.3 loading early after *de novo* infection and consequences of latency establishment in the absence of both chaperone proteins.

Our studies revealed that both HIRA and DAXX histone chaperones associate with LANA speckles independent of PML/ND10 suggesting that incorporation of these histone chaperones into LANA speckles may support loading of H3.3 histones onto KSHV episomes and regulate viral latency. In HIRA KO cells that we characterize as having a modest defect in latency control, a redistribution of DAXX from LANA speckles was noticed with an increased nuclear diffuse pattern. This is consistent with a previous study that showed that upon lytic reactivation that DAXX is partially evicted from LANA speckles and proposed this as a mechanism to restrict loading of histone onto newly synthesized linear templates^28^. LANA has previously been shown to undergo SUMOylation that facilitates association with transcriptional repressors and that reactivation leads to deSUMOylation through disruption of interactions with SUMO ligases^46^. Our interaction data of LANA domains with DAXX did not identify any specific interaction domains; however, it would be interesting in future studies to examine how differential SUMOylation of LANA may affect the localization and dispersion of DAXX in latency and lytic replication. DAXX harbors two SUMO Interaction Motifs (SIM) that are known to be important for localizing DAXX to nuclear domains including PML/ND10^47, 48^. Specifically, SUMOylated interaction partners have been shown to stabilize DAXX retention in PML bodies^49^. Future studies aimed at deciphering the temporal and spatial localization of DAXX to LANA speckles across latency and reactivation will clarify this important protein-protein interaction and implications on epigenetic maintenance of viral episomes.

While DAXX and HIRA both load H3.3 onto viral genomes, only loss of HIRA is essential for maintaining viral latency. Our data shows that in the absence of HIRA there is a site-selective effect on H3.3 loading at the major latency locus LANA that results in loss of viral latency. These findings are supported by published studies examining HIRA on DNA virus latency^50^. Previous work looking at the role of HIRA in the context of DNA virus infection or upon transfection of plasmid DNA demonstrated that HIRA was important for packaging foreign DNA into chromatin upon introduction of these foreign DNA into cells. Additionally, these studies demonstrated that in both cell culture and *in vivo* infection models that HIRA was critical for restricting lytic gene activation via regulation of host antiviral responses and therefore had higher lytic replication in the context of HIRA knockdown. Here, we also observed that HIRA KO leads to loss of viral latency with concomitant induction of a predominantly unproductive lytic virus replication.

In profiling histone modifications (H3K37me3 and H3K4me3), we did not detect broad changes in these modifications in HIRA KO cells, however, loss of HIRA had clear effects on viral gene expression. This suggests that the transcriptional regulation in this context occurs independently of overt remodeling of chromatin and instead may reflect altered H3.3 positioning, changes in transcription factor occupancy, RNA polymerase dynamics or other chromatin modifying events not captured in our analyses. Additionally, unlike previous work which showed reduced H3.3 loading on host or viral genomes with HIRA knockdown^50, 51^, our findings had the unexpected consequence of increased loading, however, specifically at the locus for LANA. This could be due to a direct effect of HIRA loss where DAXX is now able to freely load H3.3 at this locus or alternatively an indirect effect on regulation of host or viral factors regulating chromatin dynamics at the major latency locus. Future mechanistic work focused on the direct vs. indirect effects of HIRA on host and KSHV episomes in latency will help resolve the pathway in which loss of HIRA disrupts latency.

Taken together, our findings support a model where the H3.3 histone chaperones DAXX and HIRA both participate as machinery in loading H3.3 onto viral episomes. We demonstrate that H3.3 is loaded early after de novo infection (within hours) and is stably maintained on latent episomes, implicating H3.3 as both an early and persistent determinant of viral latency. While we demonstrate that both DAXX and HIRA localize to LANA speckles independent of PML bodies, HIRA has a non-redundant role in maintaining latency via site-selective regulation of H3.3 deposition within the LANA locus, suggesting that HIRA acts as a critical checkpoint for controlling latency via precise spatial H3.3 loading not global H3.3 abundance. Future studies examining how LANA coordinates viral and host pathways to maintain latency will further the understanding of herpesvirus exploitation of host epigenetic machinery to balance viral latency and reactivation.

## Materials and Methods

### Cell lines, media, antibodies and virus

BCBL-1 PEL cells were grown in complete RPMI 1640 media^52^. BCBL-1 cells overexpressing histone H3.3 were created by transducing the plasmid pOZ-HA-H3.1 or -H3.3 which couples the expression of histones with the interleukin-2 receptor chain α (IL2Rα), which was used to sort transduced cells. This system, which has been widely used to characterize genome-wide H3.3 occupancy, has also been used successfully to study HSV-1 chromatinization during lytic replication^53^.

Antibodies used were as follows: DAXX 677B rabbit (in house); DAXX 5.14 mouse monoclonal^54^; PML 14 rabbit^55^; and HIRA WC119.2H11 mouse monoclonal (generous gift from Dr. Peter Adams, Sanford Burnham Prebys Medical Discovery Institute). Antibodies for ChIP were control rabbit IgG (Santa Cruz Biotechnology, sc-2027), rat IgG (Santa Cruz Biotechnology, sc-2026), rabbit against H3K4me3 (Abcam, ab8580), H3K27me3 (Upstate, 07-449), rat LANA monoclonal antibody (ABI Inc., LN53), and rabbit anti-Histone H3.3 (phospho S31) antibody (Abcam, ab92628). Virus was produced from iSLK cells as described previously and were used to produce .219 and BAC16 virus ^56–58^.

### *De novo* infections and creation of stably infected cells

For *de novo* infections in kinetic experiments, cells were infected with BAC16 virus (300 virus particles per cell) for 2 hours and then unbound virus removed by PBS washing. Cells were then collected immediately or at post-infection time points as stated and processed for ChIP.

For creating stably infected cell lines, cells were infected as described above and selected for greater than two weeks to ensure stable maintenance of BAC16 or .219. The resulting cells were maintained in hygromycin- or puro-containing media for BAC16 and .219 respectively.

### Coimmunoprecipitation

For coimmunoprecipitation studies, 293T cells were transfected with plasmid constructs using ExtremeGene9 (Roche). After 48 hours cells were lysed in lysis buffer (7.6mM NaH_2_PO_4_, 12.4mM Na_2_HPO_4_, 250mM NaCl, 30mM NaPPi, 5mM EDTA, 10mM NaF, 0.1% NP-40, pH 7.0, complete protease inhibitor cocktail (Roche)) and quantified using BCA assay. For immunoprecipitation, 1mg of protein was incubated overnight with FLAG M2 magnetic beads (Sigma) diluted in IP buffer (50mM Tris-HCl pH 7.4, 150mM NaCl, 1mM EDTA, 1% Triton-X, complete protease inhibitor cocktail (Roche)). The following day beads were washed with IP buffer and eluted with FLAG peptide (Sigma).

### Western blot

Cells were lysed directly in Laemmli buffer. Protein samples were separated by 4-20% SDS-PAGE gels (Biorad), transferred to nitrocellulose membranes (Whatman, Dassel, Germany), and blocked with 5% non-fat milk/PBS, 0.1% Tween (PBST). Primary antibodies were diluted in 5% milk/PBST and incubated overnight at 4°C. Membranes were then washed 3X with PBST and incubated for 1 hour at RT with appropriate secondary antibody at 1:5000. Membranes were then washed 3X with PBST and were visualized by enhanced chemiluminescence (ECL).

### Immunofluorescence (IFA)

IFAs were performed as previously described with a few modifications for adherent cells^59^. In brief, cells were grown on glass cover slips in a 24-well plate. Prior to staining cells were washed twice with PBS, fixed with 1% formaldehyde, and permeabilized with PBS 0.2% Triton X-100 buffer. After two more washes, appropriately diluted primary antibody was added and incubated at room temperature for 60 minutes, followed by PBS washes and addition of a fluorescently tagged secondary antibody. Secondary antibody was washed off using PBS and coverslips were mounted on glass slides with media containing DAPI for analysis by microscopy.

### Flow cytometry

KSHV BAC16 infected HEp-2 cell supernatant was used for spin-inoculation of 293T supplemented with 10μg/mL polybrene at 2,500rpm for 90 minutes. The next morning virus containing media was removed and after 48 hours cells were fixed, washed in PBS, and resuspended in FACS Buffer (PBS, 2% FBS, 1mM EDTA). Cells were analyzed on the BD Symphony A3 with gating done on uninfected 293T (GFP^-^) and 293T-BAC16 (GFP^+^). Data were analyzed in FlowJo.

### Production of HEp-2 cells with FLAG-tagged H3F3B gene, DAXX and HIRA knockouts

HEp-2 cells were electroporated using the Neon Transfection system (Invitrogen) with equimolar amount of the crRNA/tracrRNA (Integrated DNA Technologies) targeting 5’-terminus of the H3F3B gene, the single-stranded oligonucleotide donor template containing the FLAG sequence and the Alt-R S.p. Cas9 nuclease (Integrated DNA Technologies). After 3 days cells were seeded 1 cell/well into 96-well plates for clonal growth. Clones were screened by immunofluorescence with FLAG antibodies. The correct knock-in of the FLAG tag into H3F3B gene was confirmed by DNA sequencing. DAXX and HIRA knockouts in HEp-2 cells were produced as previously described^51^.

### mRNA analysis using qPCR

WT BAC16 virus infected control, HIRA KO, and DAXX KO cells were seeded in a 6-well plate in triplicate. The cells either remained untreated or were treated with inducer (2 mM sodium butyrate) and were incubated at 37°C for 48 hours. The cells were washed with PBS, and RNA was isolated by using the previously described RNA-Bee protocol^60^. DNase digestion of RNA was performed using the Turbo DNase kit from Ambion (AM 2238). RNA was then converted to cDNA using the ABI High-Capacity RNA to cDNA kit. The cDNA was analyzed using the ABI StepOne or Roche LightCycler System for the expression of the latent gene LANA, lytic switch gene RTA, and the late genes ORF19 and ORF59 using SYBR green chemistry.

### Chromatin Immunoprecipitation (ChIP)

ChIP-seq was performed to assess LANA, H3.1 and H3.3 binding on the KSHV genome in BCBL-1 cells, and to assay H3K4me3 (activating) and H3K27me3 (repressive) epigenetic marks. The kinetics of H3.3 deposition on the KSHV epigenome post de novo infection of SLK cells were measured by ChIP-qPCR. For ChIP-seq, 10 million cells and for ChIP qPCR following de novo infection, 100,000 cells per antibody were used in triplicate for chromatin preparation. Cells were washed with PBS (room temperature), fixed with 1% formaldehyde in the plate for 10 min at room temperature followed by quenching of formaldehyde by adding 2.5 M glycine to a final concentration of 0.125 M. Cells were washed twice with PBS, and 10 mL ice cold Farnham lysis buffer (5 mM PIPES [pH 8.0], 85 mM KCl, 0.5% NP-40) with protease inhibitors was added to the plate. Cells were scraped off mechanically and centrifuged at 3000 rpm for 5 min at 4°C. Cells were further incubated in Farnham lysis buffer on a rocking platform for 5 min and were pelleted. The nuclei thus obtained were resuspended in 160 μL RIPA buffer (1% NP-40, 0.5% sodium deoxycholate, in 1×PBS with protease inhibitor). For 293T cells, 0.3% SDS was used, and for iSLK cells 0.7% SDS. The chromatin was sheared for a peak enrichment of approximately 300 bp in a Diagenode Bioruptor for SLK with settings power output H; three times 5 min each (0.5% SDS RIPA) and one-time M output for 5 min. For BCBL-1 cells (0.3% SDS RIPA) Power output H; two times 5 min each. The chromatin was obtained by centrifugation of the cells at 14000 RPM for 15 min at 4°C. The obtained chromatin was diluted with the RIPA buffer to adjust the final SDS concentration to 0.1%. Antibodies (5 µg/10 million cells) of interest were subsequently added. Antibody and chromatin complex was incubated overnight at 4°C.

The complex was then subjected to binding with pre-washed 20 µl Pierce protein A/G magnetic beads (88802) (PBS 10% BSA). The bead, antibody and chromatin complex was incubated in cold room for another two hours. Magnetic separation was performed to collect the beads. Beads were washed five times with 1 mL LiCl wash buffer with protease inhibitor and once with TE. The DNA was reverse-crosslinked by incubating twice in 150 μL elution buffer. DNA was isolated by using Monarch PCR purification kit and resuspending in 50 μL water. For qPCR, 1 μL was used per 10 μL reaction and either IgG or input was used for quantification and control.

### ChIP Sequencing

ChIP libraries were created using NEB Ultra II kit (E7645S) for Illumina as per the manufacturer’s recommendations. The sequencing was performed by multiplexing the libraries on HiSeq 3000 platform using 1*150 runs at the University of Florida ICBR.

The subtle differences in chromatin modifications were observed by performing normalization strategies using Drosophila chromatin spike-in strategies with some modifications^61^. We utilized Active Motif “spike-in” control chromatin and antibody (61686 and 53083) kits. Before performing the ChIP, the chromatin from 10 million cells was pre-mixed with 50 ng “spike-in” control chromatin made from Schneider’s Drosophila Line 2 (S2) cells and 2 µg per 10 million cells of Drosophila-specific histone variant, H2Av antibody. 4 µg specific antibody of interest was then added to perform ChIP as described above.

The ChIP-seq was analyzed as follows. Paired-end reads were quality checked using FastQC. Adaptors were trimmed using the Trimmomatic program with the settings, ILLUMINACLIP: adapters.fa:2:30:10, LEADING:3, TRAILING:3 and MINLEN:36. Trimmed reads were independently aligned to the KSHV viral genome (NCBI:GQ994935), the Drosophila genome (dm6), and the human genome (hg19) using bowtie2 retaining only the best alignment for every read. The mapped reads were sorted using samtools and duplicate reads were removed using Picard MarkDuplicates. For the viral and human alignments, ChIP-seq peaks were called using MACS2 with the setting --nomodel and --extsize 147 and without specifying any control samples. To quantitatively scale the peaks on the viral genome, the scaling factor was calculated based on the number of reads that aligned to the spike-in (Drosophila) sample as previously described^61^. To account for differences in viral episome copies between infected WT, DAXX KO, and HIRA KO cells, we calculated an ’episome factor’ as the ratio of number of aligned reads in input from control cells (WT) to input from knockout cells (DAXX KO or HIRA KO). The effective scaling factor was the product of spike-in scaling factor and the episome factor. In case of human samples, only the spike-in factor was used as the effective scaling factor. The peaks were scaled using a custom python script. All scaled and unscaled bedgraph (BDG) files were then converted to wiggle (WIG) files for visualization in IGV.

### Cleavage Under Targets & Release Using Nuclease (CUT&RUN)

CUT&RUN was used to map FLAG-H3.3 in HEp-2 cells following the Active Motif kit (53180). This protocol leverages the fusion of protein A to micrococcal nuclease (pA-MNase) to selectively cleave antibody-bound chromatin in nuclei of cells that are immobilized on concanavalin (conA) beads. Cleaved fragments are released into the supernatant and collected for library construction and Illumina Sequencing. Antibodies for FLAG (Biolegend) & IgG (Biolegend) were individually added to each sample. Cleaved fragments were library prepped using the NEB Ultra II library preparation kit for Illumina sequencing libraries.

### CUT&RUN Data Analysis

Samples were prepared as 150-bp, paired-end barcoded libraries and sequenced on the Illumina NovaSeq 6000 instrument at the University of Florida Interdisciplinary Center for Biotechnology Research (ICBR) NextGen DNA Sequencing Core. Sample quality was analyzed by FastQC and adapter trimming and quality processing by Trimmomatic. Reads were aligned with Bowtie2 to a concatenated, repeat-masked genome of human GRCh38/hg38 and KSHV BAC16 genome (GQ994935.1). Aligned BAM files were used to generate KSHV genome tracks visualized in IGV.

Initial analyses of gene tracks for samples indicated a selection for smaller genomes. To normalize for this, samples were scaled based on a scale factor created from sequenced input. Briefly, coverage from input samples were binned (500bp) across the genome to examine read count depth on the left and right side of genome. A scale factor was then generated, and the left end of genomes were scaled to account for enrichment of smaller episomes consistent on the right end of genomes. Scaled KSHV genome tracks were used for FLAG H3.3 visualization in IGV.

## Data Availability Statement

All data presented here are available from Gene Expression Omnibus (GEO) under the accession number GSE224343. The accession number will be incorporated in the manuscript when it is available.

## Acknowledgements

This project was funded by NCI/P01CA214091 to R.R. and NIDCR R01 DE026707 to R.R./A.I. and support of UFHCI to R.R./A.I. S.M. was supported by NCI T32CA257923. The authors would like to thank the University of Florida Interdisciplinary Center for Biotechnology Research (UF-ICBR) Cytometry Core (RRID:SCR_019119) and Gene Expression and Genotyping Core (RRID: SCR_019145) for their expertise.

---

**Supplementary Figure 1. Validation of H3.3 loading in BCBL-1.** (A) BCBL-1 cells were transduced with HA-H3.1- or HA-H3.3-expressing retrovirus and sorted with IL2Ra antibody-conjugated magnetic beads. Lysates were analyzed by western blotting with anti-HA mAb and GAPDH loading control. (B) ChIP-qPCR in BCBL-1 to examine H3.3 loading (H3.3-S31P) at viral promoters of LANA, K7, and ORF19.

**Supplementary Figure 2. Interaction of H3.3 chaperones with LANA deletion constructs.** (A) Schematic of N-terminally FLAG-tagged LANA domain deletions used for coimmunoprecipitation assays. (B) Coimmunoprecipitation assays in 293T of FLAG-tagged LANA domain deletions with HA-tagged HIRA or GFP-tagged DAXX constructs. Expression of constructs is observed from input samples in IP assays.

**Supplementary Figure 3. Characterization of DAXX and HIRA knockout and H3.3-FLAG knock-in HEp-2 cells.** (A) CRISPR/Cas9 was used to produce DAXX KO and HIRA KO HEp-2 cells with a FLAG knock-in into the endogenous H3.3 locus. The FLAG-encoding sequence was introduced at 5’ end of the H3F3B gene by homologous recombination. Single-cell clones were characterized for DAXX and HIRA KO and expression of FLAG-H3.3 using western blot. In control (CTL), HIRA KO, and DAXX KO cells, several clones showing FLAG-H3.3 expression were obtained. WT HEp-2 were used as control. Actin was used as loading control. (B) Microscopy analysis of endogenous tagged FLAG-H3.3. Top: exponentially grown cells; middle: metaphase spread confirming chromosome association of FLAG-H3.3. FLAG (green): DNA (blue). Bottom: Distribution of H3.3 in KSHV infected HEp-2 cells. H3.3 (FLAG, green) has homogenous nuclear distribution in KSHV-infected (LANA, red) HEp-2 cells. DNA: blue.

**Supplementary Figure 4. Normalization of FLAG H3.3 signal in CUT&RUN data.** (A) Gene tracks of FLAG H3.3 signal along KSHV genome in control, DAXX KO, and HIRA KO HEp-2 cells by CUT&RUN (B) Distribution of counts by 500bp bins of KSHV genome for CTL, DAXX KO, and HIRA KO cells. Signal breakpoint for smaller episomes was identified and a scale factor was produced as the ratio of the right side of genome to left side of genome that was applied to samples for normalization.

