## Supplementary figures and images for "Non-canonical histone H3.3 and its chaperones HIRA and DAXX participate in the regulation of KSHV latency"

### Supplementary Figure 1

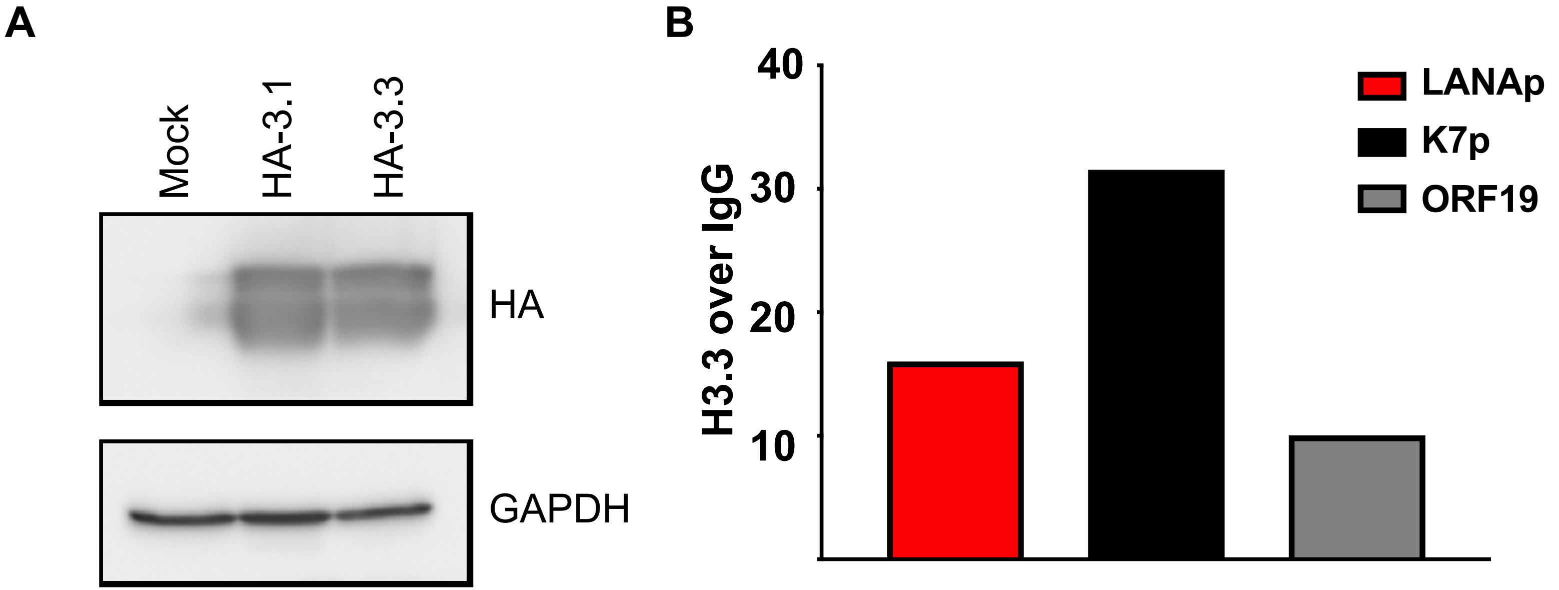

### Supplementary Figure 2

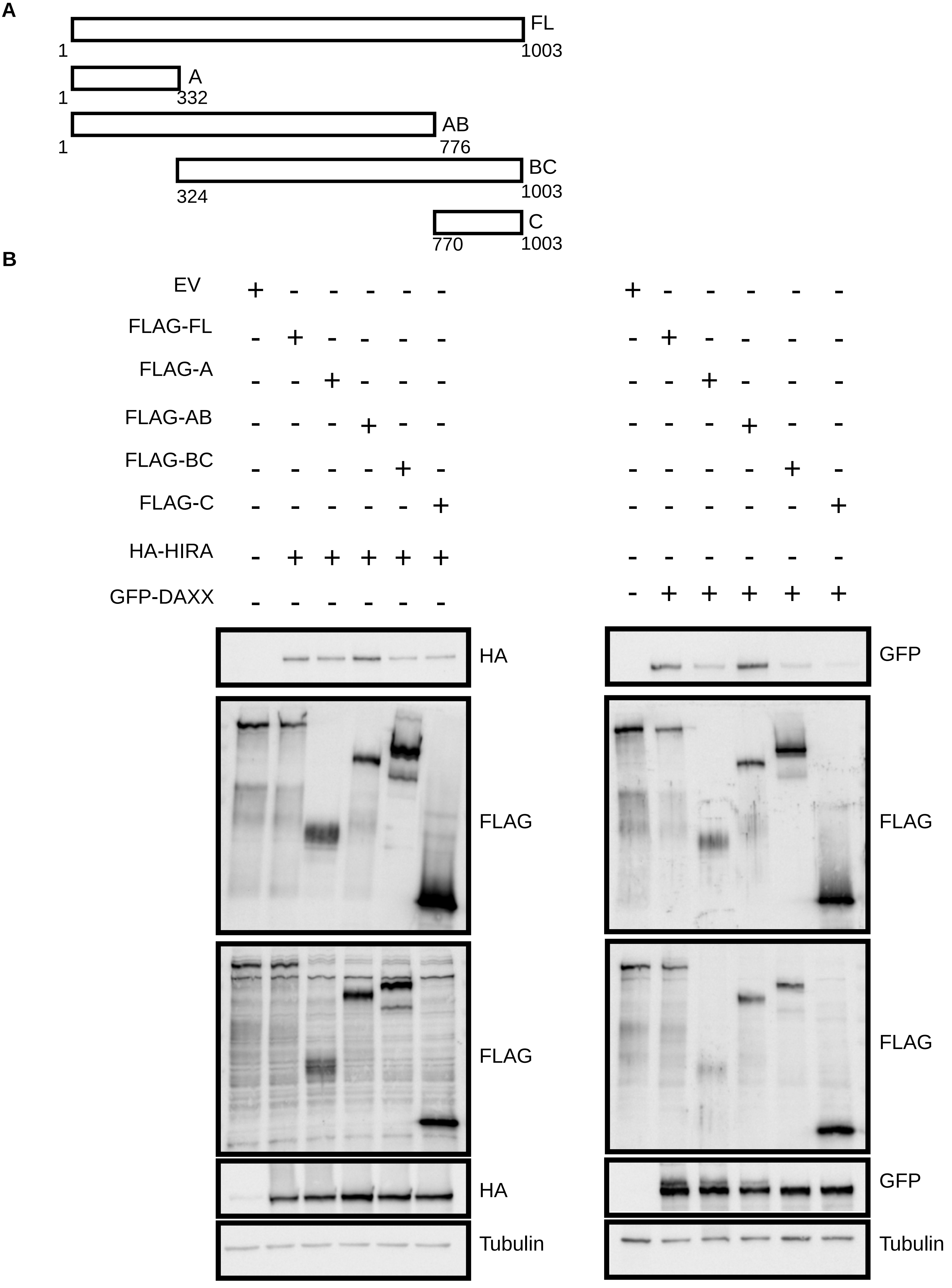

### Supplementary Figure 3

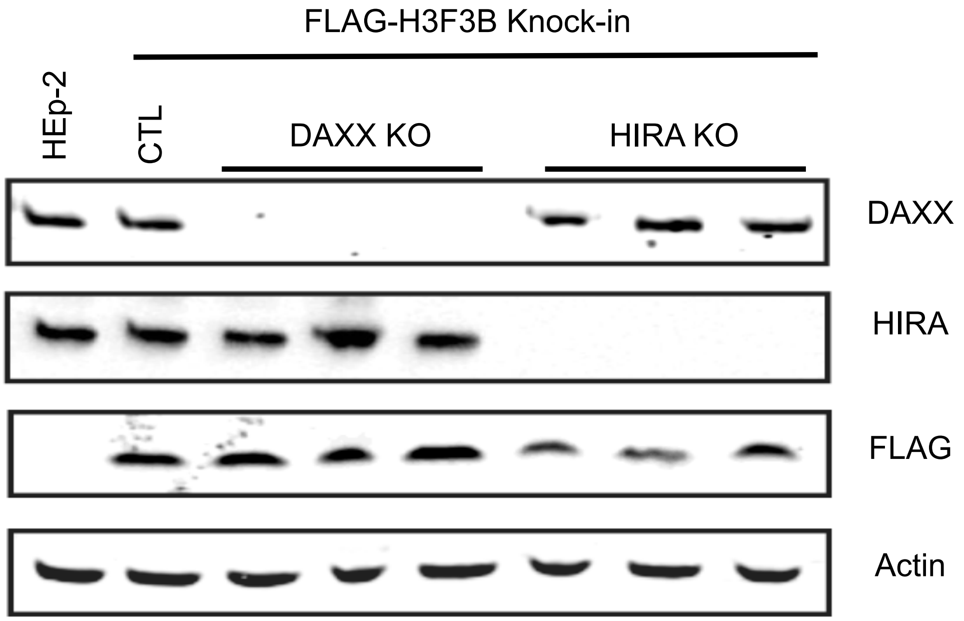

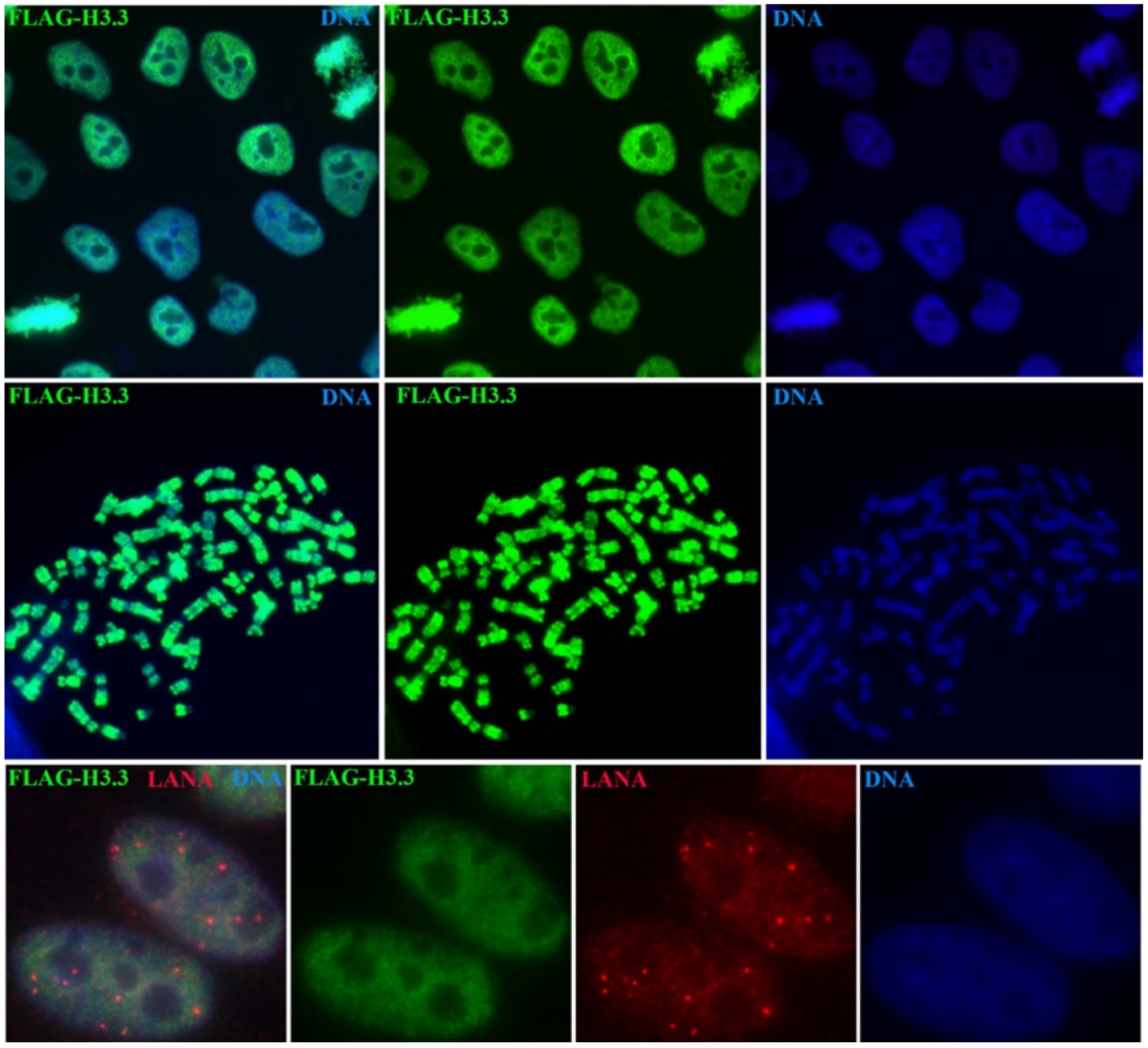

### Supplementary Figure 4

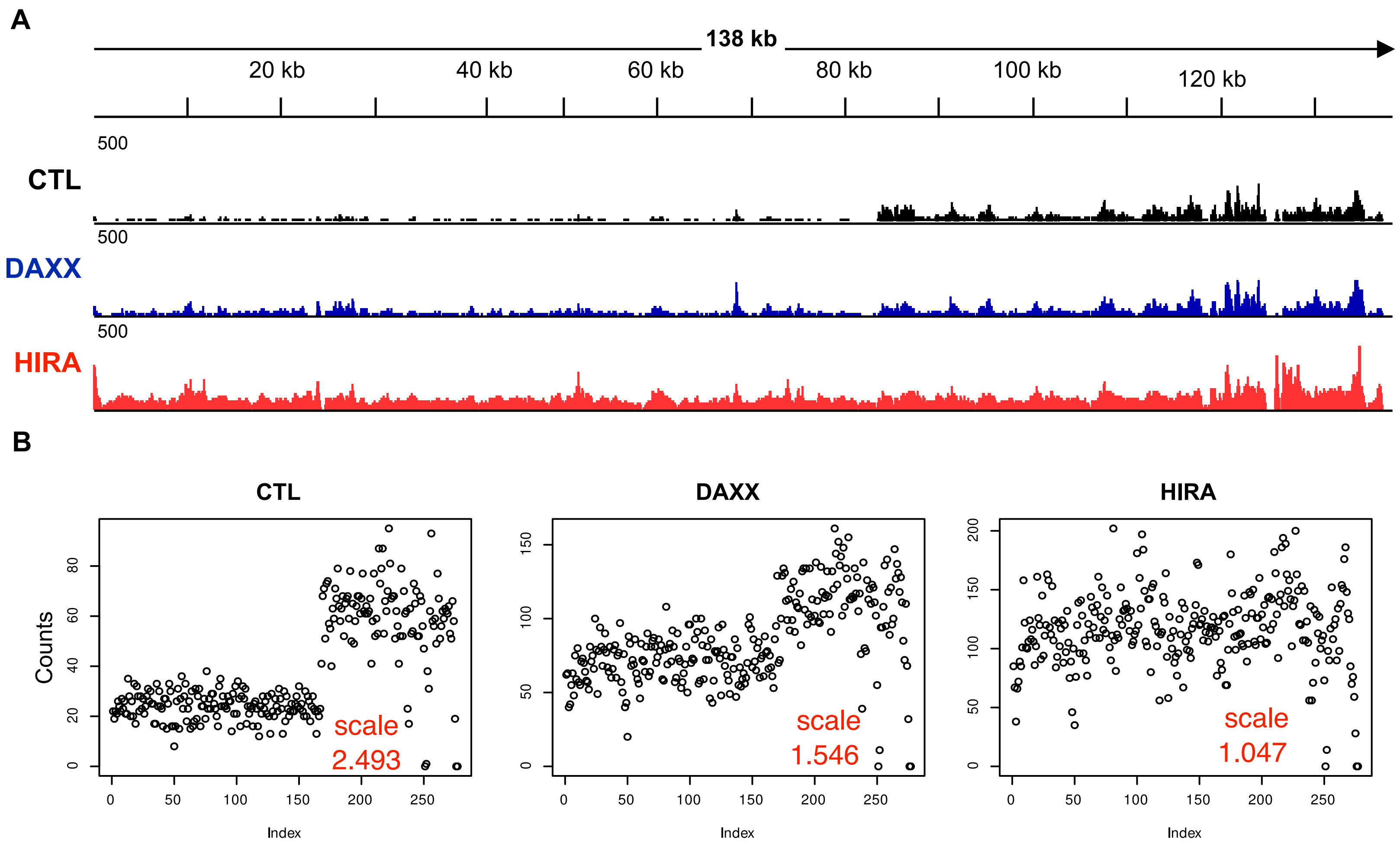
